# Differential effects of aquaporin-4 channel inhibition on BOLD fMRI and Diffusion fMRI responses in rat visual cortex

**DOI:** 10.1101/2020.01.24.917849

**Authors:** Yuji Komaki, Clément Debacker, Boucif Djemai, Luisa Ciobanu, Tomokazu Tsurugizawa, Denis Le Bihan

## Abstract

The contribution of astrocytes to the BOLD fMRI and DfMRI responses in visual cortex of mice following visual stimulation were investigated an aquaporin 4 (AQP4) channel blocker, TGN-020, acting as an astrocyte function perturbator. Under TGN-020 injection the amplitude of the BOLD fMRI response became significantly higher. In contrast no significant changes in the DfMRI responses and the electrophysiological responses were observed. Those results further confirm the implications of astrocytes in the neurovascular coupling mechanism underlying BOLD fMRI, while DfMRI relies on other, not astrocyte-mediated mechanisms.

## Introduction

Diffusion functional MRI (DfMRI) has been proposed as an alternative to blood oxygenation level dependent (BOLD) fMRI to monitor neural activity noninvasively [1]. Several studies have demonstrated that that DfMRI and BOLD fMRI responses to a variety of stimuli differed qualitatively and quantitatively (ie amplitudes and time courses of responses) [1–5] suggesting that mechanisms underlying BOLD and diffusion fMRI must be different, although this view has been controversial [6,7] While BOLD fMRI relies on the indirect neurovascular coupling mechanism [8,9] the current hypothetical mechanism of DfMRI is thought to be related to the neural activation triggered cell swelling for which there is a large body of evidence [10,11]. Beside the established fact that water diffusion as monitored with MRI decreases in tissues undergoing cell swelling in pathological, extraphysiological and physiological conditions [12–16] several preclinical studies relying on pharmacological challenges interfering with neurovascular coupling or cell swelling have confirmed that 1/ the DfMRI and BOLD fMRI responses could be decoupled, confirming their differential mechanisms; 2/ the DfMRI response is not dependent on neurovascular coupling, but, instead, sensitive to underlying neural swelling status [17–19] and 3/ the DfMRI response follows neural activity status closely and more accurately than BOLD fMRI, especially under anesthetic or vasoactive drug conditions [17,20].

To further uncover the differences between DfMRI and BOLD fMRI mechanisms we investigated the contribution of astrocytes to both responses. Astrocytes have been shown to play a major role in the neurovascular coupling mechanism [21]. Interfering with astrocyte function should, thus, impact BOLD fMRI responses, but not necessarily neural responses which have been shown to persist unaltered after neurovascular coupling inhibition [19]. Hence, DfMRI responses should remain relatively immune to astrocyte activity if they originate directly from neurons. To test this hypothesis, we have used an aquaporin-4 channel blocker (2-(nicotinamide)-1,3,4-thia-diazole, TGN-020) [22]. In the brain aquaporin-4 (AQP) channels are exquisitely expressed on astrocytes cell membranes. mainly at the astrocyte end-feet surrounding vessels in the perivascular spaces, regulating water flow between blood and brain [23] and, in turn, the astrocyte volume and cerebral blood flow (CBF) [24].

## Material and methods

### Animals

The study was performed on 34 adult mice (20-28g, C57BL/6J, male, Charles River laboratories, Lyon, France). Mice were housed in groups of six under a 12-hour light/dark cycle, with access to food and water *ad libitum*.

All animal experimental procedures were performed in accordance with the EU Directive 2010/63/EU for care and use of laboratory animals and approved by the Comité d’Ethique en Experimentation Animale (CETEA) de la Direction des Sciences du Vivant (DSV) du Commissariat à l’Energie Atomique et aux Energies Alternatives (approval number: APAFIS#8462-20170109l5542l22 v2).

### Functional MRI acquisitions

MRI was conducted using a 17.2 Tesla MRI system (Bruker BioSpin, Etlingen, Germany) with a 25mm quadrature birdcage coil (RAPID Biomedical GmbH, Rimpar, Germany). The mice were anesthetized with isoflurane (1-1.5% in medical air containing 30% O_2_) and placed inside the magnet in a dedicated animal bed. The respiratory cycle and body temperature were monitored during scanning (model 1025, SA Instruments, NY, USA). The body temperature was maintained at 37 °C by means of circulating hot water. An optical fiber for light stimulation was placed in front of the right eye of the mouse. After completing the animal set-up, the anesthesia was switched from isoflurane to medetomidine (s.c. 0.1 mg/kg bolus, 0.2 mg/kg/h continuous infusion, Orion Pharma, Espoo, Finland). The functional MRI acquisition started 30 minutes after the administration of the medetomidine bolus.

High resolution anatomical images of the whole brain were acquired using a Rapid Acquisition with Relaxation Enhancement (RARE) sequence with the following parameters: effective echo time (eTE), 23 ms; repetition time (TR), 2000 ms; RARE factor, 8; number of averages, 4; spatial resolution, 75 x 75 x 500 (μm)^3^; number of slices, 21.

Functional BOLD and diffusion fMRI acquisitions were performed using a double spin echo echo-planar imaging (SE-EPI) sequence to mitigate the effects of eddy currents and background magnetic field gradients: TE, 24.5 ms; TR, 2000ms; number of averages, 1; spatial resolution, 200 x 200 x 1000 (μm)^3^; number of slices, 9; b-values, 0 (BOLD), 1000, 1800 s/mm^2^; number of repetitions, 180.

The visual stimulus consisted in six blocks of a blue light (20s, 2Hz, 10ms pulse duration) alternating with darkness rest periods (40s) using a light emitting diode (LED) and Arduino programming board (ArduinoCham, Switzerland). The Arduino programming board synchronized the trigger from the Bruker scanner during the fMRI scanning. BOLD fMRI data were acquired twice, DfMRI three times with b = 1000, and six times with b = 1800 in each session (before and after TGN-020 administration). TGN-020 (200 mg/kg i.p., Merck KGaA, Darmstadt, Germany), an AQP4 inhibitor [22], was administrated after an initial set of baseline BOLD fMRI and DfMRI measurements. Functional data were collected again after 15 minutes from TGN-020 administration. TGN-020 was administered to a group of nine mice, while the other nine (control group) received a saline injection.

### MRI data analysis

As previously described [25], SPM12 software (Welcome Trust Center for Neuroimaging, UK) and a tailor-made program (Matlab, The MathWorks, Inc., USA) were used for fMRI analysis. Image processing, consisting of slice timing correction, motion correction, normalization of brain coordinates, and smoothing (Gaussian kernel with FWHM of 0.6 mm), was performed for all fMRI data before statistical analysis. Statistical t-maps were calculated using a generalized linear model. Activation was detected using a statistical threshold of p < 0.05 (false discovery rate (FDR) corrected for multiple comparisons). Activities before and after administration were compared using paired t-test (p<0.05 FDR corrected). Region of interest (ROI) of primary visual cortex (V1) for the time course analysis were defined using the Allen Mouse Brain Atlas [26,27]. The signal change was expressed as percentage with the average value at rest taken as 100%. Time courses before and after administration were compared using paired t-tests.

The Apparent Diffusion Coefficient (ADC) (in mm^2^/s) was obtained at each time point using the following equation:

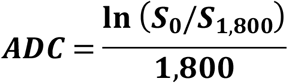

where S_0_ and S_1800_ are signal intensities obtained for b = 0 and 1 800 s/mm^2^, respectively.

### Local field potentials (LFP) recordings

Electrophysiological recordings were performed separately, outside the MRI bore. The animals, first anesthetized with 1.5% isoflurane, were placed in a stereotaxic frame (David Kopf Instruments, CA). The body temperature was maintained at 37°C using a heating pad (DC temperature controller; FHC Inc., Bowdoin, ME, USA). The skull was exposed and multiple holes (1 mm diameter) were made with a dental drill for insertion of micro-electrodes. The multiple tungsten microelectrodes (< 1.0 MΩ, with 1 μm tip and 0.127-mm shaft diameter, Alpha Omega Engineering, Nazareth, Israel) were positioned on the left visual cortex (AP −3.5 mm, ML −2.2 mm, DV −1.5 mm from the Bregma). After surgery, the anesthesia was switched from isoflurane to medetomidine (s.c. 0.1 mg/kg bolus, 0.2 mg/kg/h continuous infusion, Orion Pharma, Espoo, Finland) as in the fMRI protocol. Electrodes were connected to a differential AC amplifier Model 1700 (AM systems, Sequim, WA, USA), via a Model 1700 head stage (AM systems, Sequim, WA, USA). The LFP and MUA signals were acquired at 10 kHz sampling rate using dedicated data acquisition software (Power Lab, AD Instruments, Dunedin, New Zealand). The reference electrode was positioned on the scalp. The visual stimulation paradigm was the same as for fMRI, with six blocks of alternating blue light stimulation (20s, 2Hz, 10ms pulse duration) and rest (40s) periods in a dark room.

### LFP analysis

The raw electrophysiological signals were frequency-filtered with less than 100 Hz for LFP [28]. The power of LFP was calculated at each time point. The AUC during the stimulation period (10-20 s) was compared with the pre-stimulation period (0-10 s). The peak amplitude was defined as the maximum value during stimulation.

## Results

### Local Field Potentials

LFP amplitude increased in V1 upon 2Hz blue light stimulation (fig. 1a, b). There was no difference in the area under the curve (AUC) and the peak amplitude of LFP responses after saline or TGN 020 administration (p<0.05) (Fig. 1c, d) confirming that interference on astrocyte function induced by TGN 020 has no effect on neuronal responses in V1.

**Figure 1:**
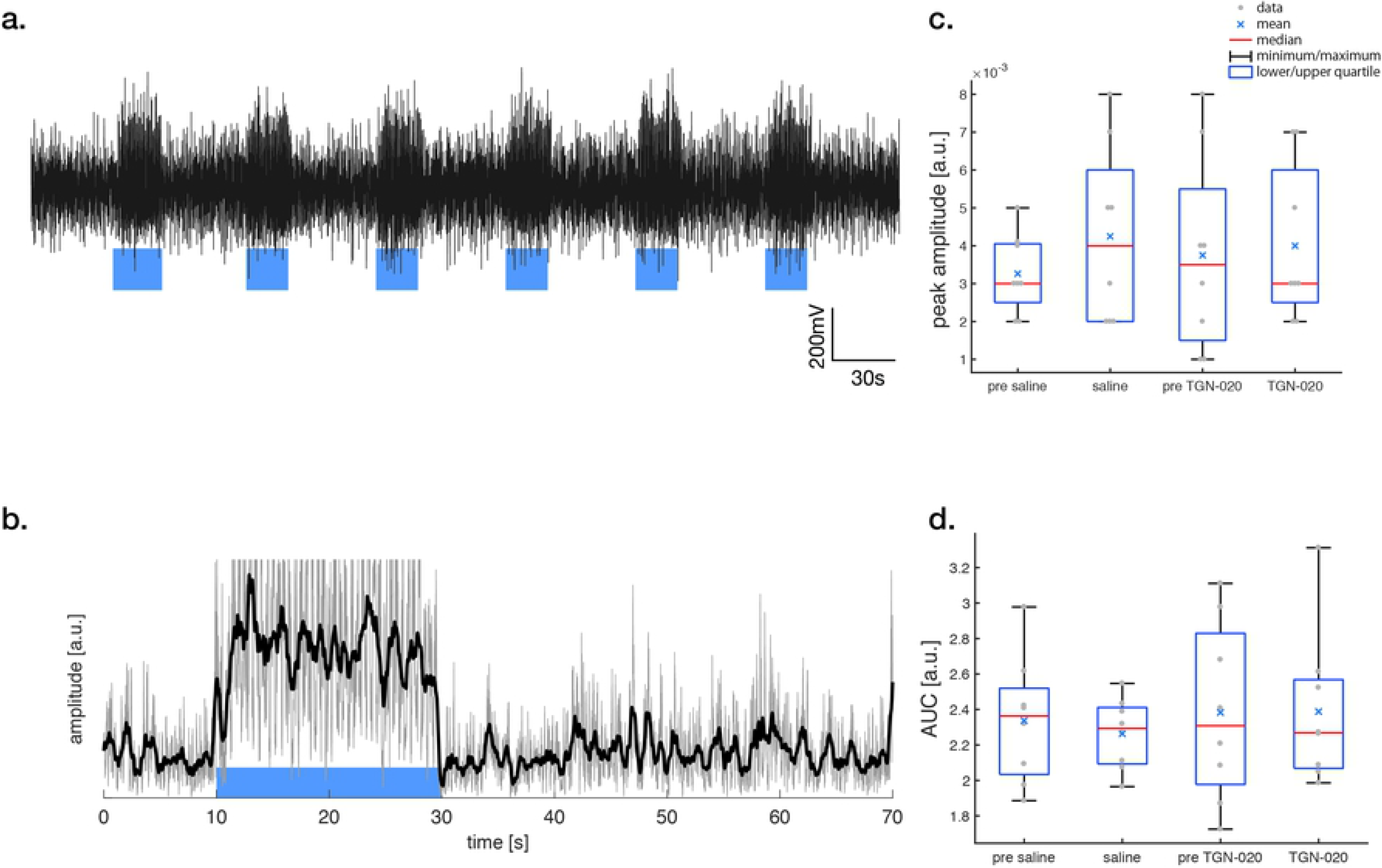
LFPs. Local Field Potential (LFP) in V1 under blue light stimulation (a). The visual stimulus was applied between 10 and 30 seconds (a, blue block). A 100 Hz low pass filter was applied to the acquired data (b, gray line). Signal obtained by applying a moving average filter of 0.5 s window width (b black line). The boxplot of peak amplitude is the maximum value during stimulation (c). The area under the curve during the stimulation period (10-30 s) was compared with the pre-stimulation period (0-10 s) (d). The difference between these data shows no significance between the conditions (saline or TGN-020) (p<0.05 Bonferroni correction with a paired t-test.).

### BOLD fMRI

BOLD activation maps (b = 0 s/mm^2^) are shown in Fig. 2a. BOLD fMRI responses of V1 were readily observed following blue light stimulation, (as well as superior colliculus, SC) and lateral geniculate nucleus, LGN). The time course of the BOLD fMRI signals in V1 are shown in Fig. 3a. While the overall time course of the BOLD fMRI responses was not different after saline or TGN-020 injection (Fig. 3a and Fig.4), their amplitudes were significantly higher (p<0.05) under TGN-020 than under saline.

**Figure 2:**
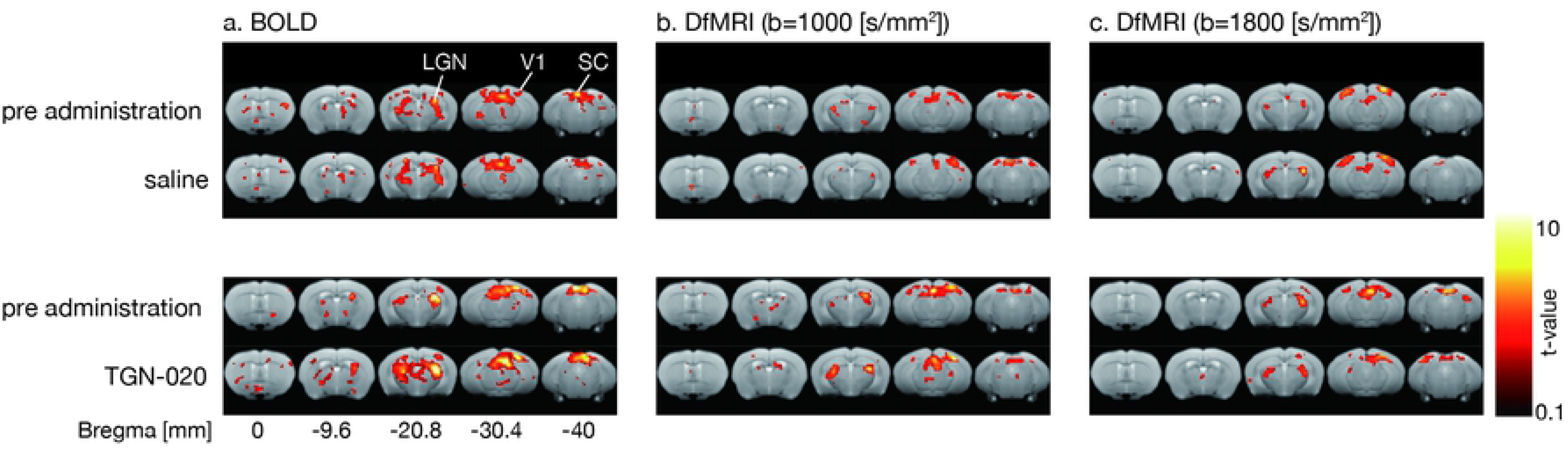
Activation maps (a: BOLD, b: DfMRI b1000; c: DfMRI b1800) **C**omparison of activation map before and after TGN-020 or saline administration (p<0.05, FDR corrected). (a) BOLD, (b) DfMRI (b=1000 [s/mm^2^]), (c) DfMRI (b=1800 [s/mm^2^]). Activation was observed in V1, SC, LGN.

**Figure 3:**
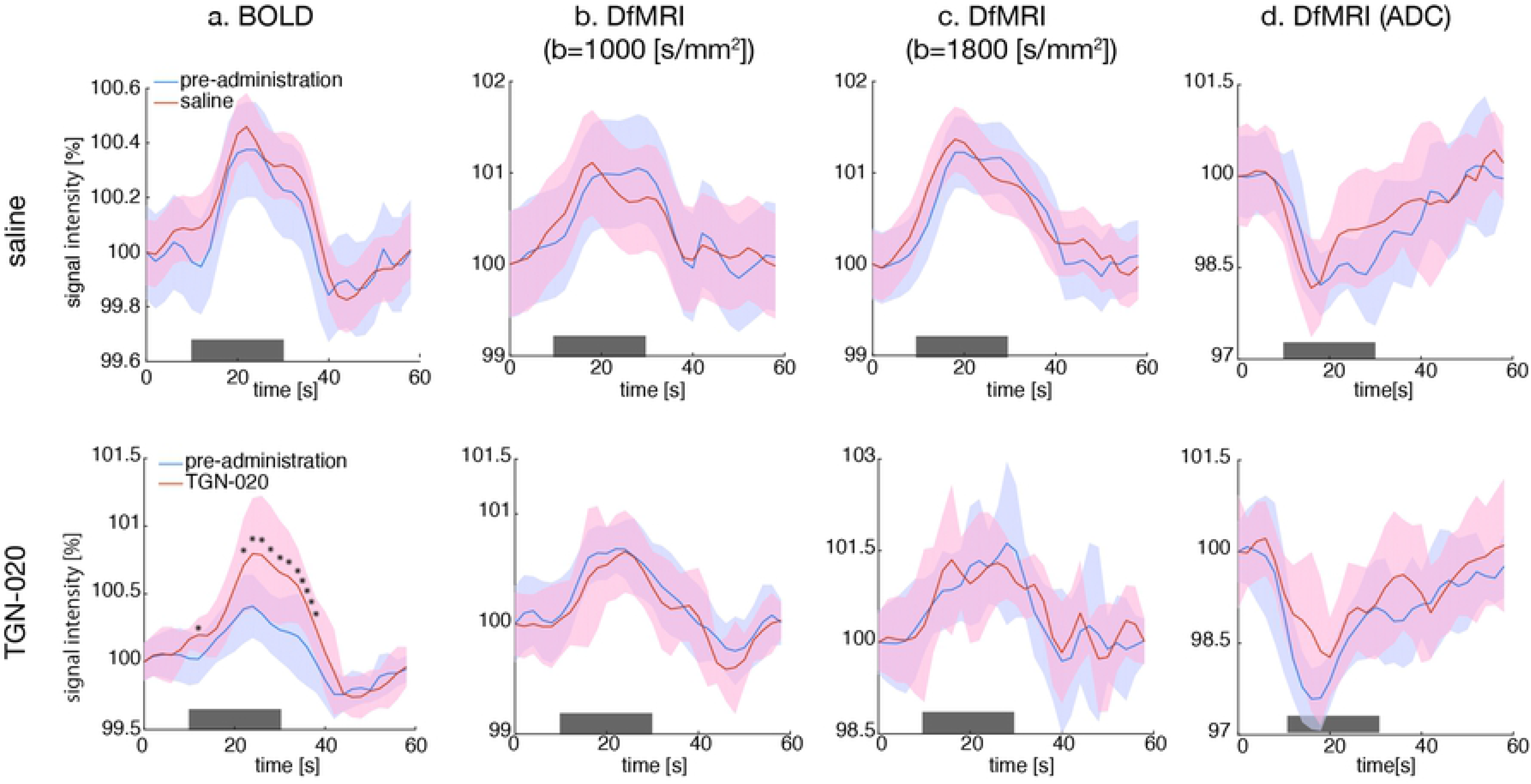
Time courses (a: BOLD; b: b1000; c: b1800; d: ADC) Time courses of signal in V1 with saline (upper row) and TGN-020 (lower row). (a) BOLD, (b) DfMRI (b=1000 [s/mm^2^]), (c) DfMRI (b=1800 [s/mm^2^]), (d) DfMRI (ADC). The amplitude of the BOLD response after TGN-020 administration (red line) is significantly larger than the pre-administration response (blue line), while the amplitude of the DfMRI and ADC response after TGN-020 administration is not significantly different. The color bands represent the standard deviation (SD) between subjects (n=9). The visual stimulus was applied between 10 and 30 seconds (gray bar). An asterisk indicates a significant difference between pre-administration and post-administration (paired t-test, p<0.05). Note that the onset and offset of the DfMRI signal response occur earlier than the BOLD fMRI responses.

**Figure 4:**
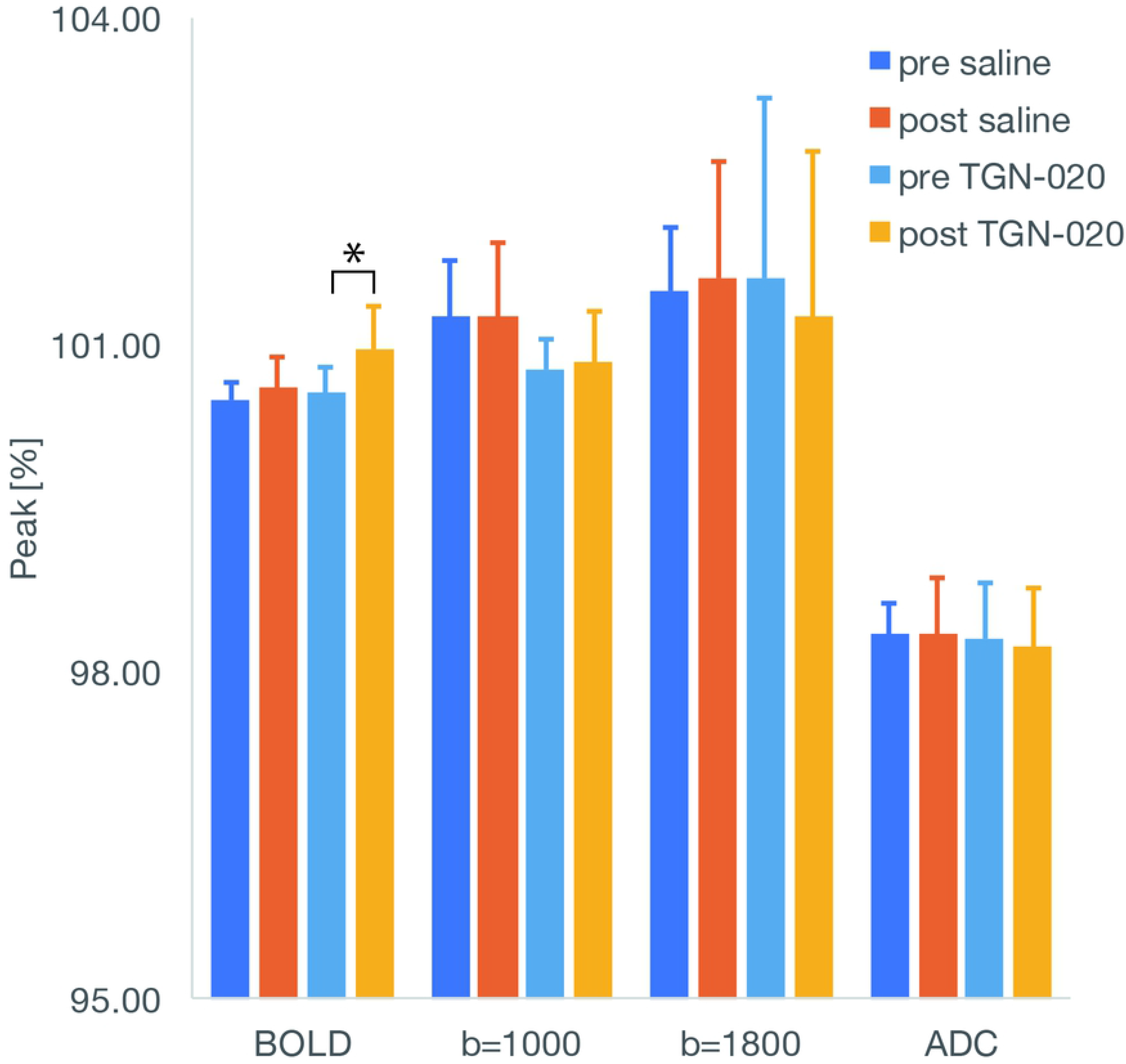
Bar plots for peak amplitude responses. Peak amplitude response before and after TGN-020 or saline administration. The peak amplitude of the BOLD signal change was significantly higher after TGN-020 administration, no change was observed for the DfMRI signal at b=1000, b=1800s/mm^2^, and for the ADC. The error bar represents the SD between subjects (n=9). An asterisk indicates a significant difference between pre-administration and TGN-020 administration (p<0.05, paired t-test).

### Diffusion fMRI

The DfMRI responses (b=1000 and b=1800s/mm^2^) were also clearly observed in V1 (as well as in SC and LGN), but with a slightly smaller spatial extent than that of BOLD fMRI (Fig.2 b,c). The amplitudes of the DfMRI responses in V1 were slightly higher than BOLD fMRI responses before saline or TGN-020 injection (Fig.3 b,c). The amplitudes remained unchanged (p<0.05) after administration of TGN-020 or saline (Fig. 3 b,c and Fig.4). The amplitude of the b1800 DfMRI response was slightly higher than the b1000 DfMRI response, corresponding to the water diffusion decrease observed upon visual stimulation, as reflected in the ADC time courses computed from b=0 and b=1800 s/mm^2^ (Fig.3d). However, the ADC change was not significantly different between saline and TGN-020 injected groups (p < 0.05) (Fig.4). The mean ADC baseline value (before stimulation) was not significantly different (p<0.05) before and after injection of saline (0.62 and 0.61 10^-3^ mm^2^/s, respectively) and before and after injection of TGN-020 (0.67 and 0.71 10^-3^ mm^2^/s, respectively).

## Discussion

BOLD fMRI has been widely used in research and clinical practice to investigate brain function noninvasively. BOLD contrast results from the magnetic susceptibility balance between oxy- and deoxy-hemoglobin in circulating erythrocytes [29] which reflects the activation induced increase in blood oxygen consumption and blood flow in cerebral blood vessels [8]. Hence, BOLD fMRI primarily reflects changes in hemodynamics and oxygenation in large and small blood vessels, not directly neural activity, resulting in known limitations, namely its limited spatial and temporal resolution with regards to underlying neural activity [9,30], its sensitivity to underlying local organizational structure of the vascular network (which may not always covariate with local neural networks [30, 31]) and to any confound interfering with the neurovascular mechanism (underlying pathology, presence of drugs) [9].

To overcome the limitations of BOLD fMRI, alternative fMRI imaging methods have been previously proposed. One of them, DfMRI, monitors changes in water diffusion occurring in activated brain tissue [1,17]. DfMRI has been shown to be more accurate in time and space than the BOLD response [5,32,33] and is thought to be more directly reflecting neural activation status. Activation-induced cell swelling has been proposed as its hypothetical mechanism [10,11] and recent studies have shown that the DfMRI signal is not related to the neurovascular coupling [19] and that, contrarily to BOLD the DfMRI signal is modulated by neuronal swelling inhibition and cell swelling facilitation [17], mirroring LFP responses. While the likely cell population involved is neuronal (dendritic spines) contribution of astrocytes has not been ruled out.

Here we used another pharmacological challenge based on the inhibition of AQP4 channels carried specifically by astrocytes. TGN-020 is known to increase astrocyte swelling, reduce water flow from astrocytes into the peri-capillary Virchow-Robin space, reduce peri-capillary fluid pressure and capillary lumen expansion and increase regional baseline CBF [34]. It has been suggested that astrocytes play a major role in neurovascular coupling [21]. Excitatory events can drive activity in interneurons [35] or astrocytes [36,37] that recruit a local hemodynamic response [38]. Upon neuronal activity through Ca^2+^ signaling, astrocytes release vasoactive substances which promote arteriolar vasodilatation and cause a CBF increase [39], but exact mechanisms are still not well understood and other studies have revealed a more complicated relationship between neuronal/glial activity and BOLD responses. Here, we found that the amplitude of the BOLD response triggered by visual stimulation was increased upon astrocyte function disruption induced by blockage of AQP4 channels with TGN-020, confirming the involvement of astrocytes in the neurovascular mechanisms underlying BOLD fMRI [40–43] as neuronal activity (as assessed from LFPs) remained unchanged. Although synaptic plasticity as well as NMDA-mediated excitatory postsynaptic currents are altered in AQP4 KO mice [44], LFPs have not been found altered by acute inhibition of the AQP4.

We might speculate that the increase in BOLD fMRI responses that we observed under TGN-020 could be linked to an increase in baseline CBF which is known to increase under administration of TGN-020 [39,45]. Indeed, functionally induced changes in CBF are thought to be proportional to the underlying baseline CBF, resulting in constant relative changes in CBF upon activation [46–49]. However, administration of acetazolamide which increases CBF by vasodilation decreases the BOLD contrast under visual stimulation [49] and other studies have suggested that activation driven changes in CBF are independent from baseline [46–48]. Those conflicting results reflect the complexity of the mechanisms involved in neurovascular coupling, which in turn influences BOLD fMRI responses. In any case it was not possible to evaluate baseline CBF from BOLD signals which are only relative and not absolute by nature.

On the other hand the present results clearly demonstrate that DfMRI responses (acquired at high b values) and resulting ADC values are not affected by the disruption of astrocyte function induced by AQP4 inhibition through TGN-020 administration, together with LFP responses, contrarily to BOLD fMRI responses, thus confirming the non-vascular nature of the DfMRI responses. Furthermore, those results show that astrocytes play a little role, if any, in the DfMRI responses, indirectly confirming their neural origin. Altogether those results also confirm that DfMRI is immune to disruptions of the neurovascular events underlying BOLD fMRI, reflecting in a more robust may neural responses which could be obtained across different physiological conditions [17].

A limitation to this study was that BOLD fMRI responses were obtained from spin-echo (SE-EPI) sequences (instead of more standard gradient-echo EPI (GE-EPI) sequences). Beside convenience and reliability (the same MRI sequence was used for both BOLD fMRI and DfMRI) SE-EPI is known to be more accurate than GE-EPI (without contamination from draining veins and large vessels) and more robust to background susceptibility artifacts. However, a drawback is that SE-EPI BOLD fMRI responses are smaller in amplitudes than with GE-EPI BOLD fMRI, so that comparison of BOLD fMRI and DfMRI responses amplitudes may be biased.

An important point to underline is that, although the same SE-EPI MRI sequence was used different behaviors were observed depending on the b values (degree of diffusion weighting) associated with the sequence. With b=0 BOLD effects were solely visible. Using a high diffusion weighting (b=1000 and 1800s/mm^2^) a completely different behavior emerged as shown here. This means that the contribution of the water diffusion effect to the signal largely predominates over the contribution of T2 which remains the same whatever the b value. Residual T2 effects are, furthermore, removed when calculating the ADC which solely reflects diffusion effects. The decrease in ADC upon neural activation, as we observed, is fully consistent with earlier reports (Le Bihan, Aso, Tsurugizawa, Abe, Nunes, etc) and generally reflects a local increase in cell size, further confirming the neural swelling hypothesis of DfMRI mechanisms. Activation driven neural swelling should be distinguished from the astrocyte swelling which may have also occurred by blocking AQP4 channels. Such astrocyte swelling could result in an small ADC decrease, but it was not observed within the conditions of this study.

Another limitation is that LFPs could be recorded only in one location, V1, following visual stimulation. Recording LFPs was obviously necessary to allow interpretation of both BOLD fMRI and DfMRI responses to disentangle potential effects of TGN-020 on neural and vascular systems. However, neurovascular coupling level, astrocytes contribution and AQP4 expression might vary across brain locations. Especially, it would be interesting to investigate in the future BOLD fMRI and DfMRI responses in areas located within cerebellum and hippocampus (CA1) which are rich in AQP4 receptors.

One may rightly question why DfMRI has not yet become popular for fMRI given the limitations of BOLD fMRI. Those limitations might be acceptable for human cognitive imaging on a coarse spatiotemporal resolution, but represent an important drawback for neuroscience applications at a finer level, as allowed with upcoming ultra-high field MRI scanners [50,51] or when using preclinical models under anesthesia. The main reason for the limited usage of DfMRI is likely technical due to the relatively higher noise level observed in long TE, spinecho based diffusion MRI compared to gradient-echo based BOLD fMRI (the amplitude of the diffusion and BOLD fMRI responses are otherwise very similar as shown in this study), which may require signal averaging over repeated fMRI sessions, a potential limitation for human studies. Another possible reason is that the putative mechanism underlying DfMRI and its direct link with neural activity (ie, neuromechanical coupling) has not yet been directly evidenced. Hopefully, this uncertainty will dissipate over time given the accumulation of studies, like this one, revealing the differential mechanisms beyond DfMRI and BOLD fMRI.

## Conclusion

Disruption of astrocyte function by blocking AQP4 channels impacts BOLD fMRI responses in V1 following visual stimulation but not DfMRI responses. This discrepancy confirms that while BOLD fMRI depends on neurovascular coupling, DfMRI relies on a different mechanism not involving astrocytes.

## Acknowledgement

This work was supported by a postdoctoral fellowship from the Uehara Memorial Foundation (Japan) and by the Louis-Jeantet Prize for Medicine from the Louis-Jeantet Foundation.

## Disclosure/Conflict of Interest

The authors have no conflicts of interest to declare.

